# Genomic features of BRDT binding sites in gene units suggest transcriptional partners and specific epigenetic landscapes to regulate transcriptional activity during spermatogenesis

**DOI:** 10.1101/822874

**Authors:** Li Wang, Iouri Chepelev, Yoon Ra Her, Marcia Manterola, Binyamin Berkovits, Kairong Cui, Keji Zhao, Debra J. Wolgemuth

## Abstract

BRDT, a member of the BET family of double bromodomain-containing proteins, is expressed uniquely in the male germ line, is essential for spermatogenesis in the mouse, and binds to acetylated transcription start sites of genes expressed in meiosis and spermiogenesis. It has thus been postulated to be a key regulator of transcription in meiotic and post-meiotic cells. To understand the function of BRDT in regulating gene expression, we characterized its genome-wide distribution, in particular the features of the BRDT binding sites within gene units, by ChIP-Seq analysis of enriched fractions of spermatocytes and spermatids. In both cell types, BRDT binding sites were mainly located in promoters, first exon, and introns of genes that are highly transcribed during meiosis and spermiogenesis. Furthermore, in promoters, BRDT binding sites overlapped with several histone modifications and histone variants associated with active transcription, and were also enriched for consensus sequences for specific transcription factors, including MYB, RFX, ETS and ELF1 in pachytene spermatocytes, and JunD, c-Jun, CRE and RFX in round spermatids. Our analysis further revealed that BRDT-bound genes play key roles in diverse biological processes that are essential for proper spermatogenesis. Taken together, our data suggest that BRDT is involved in the recruitment of different transcription factors to distinctive chromatin regions within gene units to regulate diverse downstream target genes that function in male meiosis and spermiogenesis.

## Introduction

The bromodomain is a highly conserved eukaryotic 110-amino acid long motif which binds acetylated lysine residues (Mujtaba, Zeng, & Zhou, 2007) in histones (Kanno et al., 2004) and non-histone proteins as well (Barlev et al., 2001; Gamsjaeger et al., 2011; LeRoy, Rickards, & Flint, 2008; Sinha, Faller, & Denis, 2005). Bromodomains are found in chromatin-associated factors including histone modifying enzymes (Agricola, Randall, Gaarenstroom, Dupont, & Hill, 2011; Gregory et al., 2007; Jacobson, Ladurner, King, & Tjian, 2000; Wang et al., 1997), ATP-dependent chromatin-remodeling factors (Liu, Mulholland, Fu, & Zhao, 2006; Racki et al., 2009; Thompson, 2009), and transcriptional regulators (Denis, 2010; Sankar et al., 2008; Tae et al., 2011; Tamkun, 1995; Zhou & Grummt, 2005). The BET family of bromodomain proteins are distinct in that they possess two <u>b</u>romodomains and an <u>e</u>xtra-terminal (ET) domain (Taniguchi, 2016), which is believed to be involved in protein-protein interactions (Crowe et al., 2016; Matangkasombut, Buratowski, Swilling, & Buratowski, 2000). Several other minor motifs exist among the BET proteins, but they are not conserved in all eukaryotic BET family members (B. Florence & Faller, 2001; Paillisson et al., 2007; Wu & Chiang, 2007).

All the BET proteins studied to date have been implicated in transcriptional regulation at some level. The founding member of the BET family is the *Drosophila* female sterile (1) homeotic (*fs(1)h*) gene and is the only BET gene in flies. It has been shown to regulate the transcription of the segmentation genes *Ultrabithorax* (*Ubx*) and *trithorax* (*trx*) (Huang & Dawid, 1990), as well as the genes *tailless* (*tll*) and *hückebein* (*hkb*), which govern head and gut formation (B. L. Florence & Faller, 2008). The yeast *S. cerevisiae* has two BET family proteins, BDF1 and BDF2, and both are components of the basal transcriptional machinery (Durant & Pugh, 2007; Matangkasombut et al., 2000). Although it does not have histone-modifying activity, BDF1 has been shown to be involved in depositing histone variants in chromatin (Krogan et al., 2003; Sawa et al., 2004). BDF1 has also been shown to control the transcription of several members of the U family of small nuclear RNAs (Lygerou et al., 1994).

There are four mammalian BET proteins, BRD2, BRD3, BRD4 and BRDT, and all four have been implicated in transcription regulation (B. Florence & Faller, 2001). BRD2 and BRD3 have been shown to bind to the hyperacetylated body region of actively transcribing genes and may facilitate transcriptional elongation by RNA polymerase II (Pol II) (LeRoy et al., 2008), suggesting a direct role in transcription. Further, BRD2 interacts with TATA binding protein (TBP) through its first bromodomain (BD1) (Peng et al., 2007) and with E2F through its ET domain and plays a role in modulating E2F-regulated transcription (Denis, Vaziri, Guo, & Faller, 2000; Guo, Faller, & Denis, 2000). Similarly, BRD3 binds via its BD1 to multi-acetylated GATA1, and modulates GATA1-regulated transcription (Lamonica et al., 2011). BRD4 is a regulatory component of the positive transcriptional elongation factor beta (P-TEFb) complex, and is required for P-TEFb-dependent transcription (Jang et al., 2005; Yang et al., 2005). BRD4 binding via its C-terminal domain (CTD) frees cyclin T1 (CCNT1) and cyclin dependent kinase 9 (CDK9), the major components of P-TEFb, from inhibition by the 7SK small nuclear RNA and HEXIM1 (Bisgrove, Mahmoudi, Henklein, & Verdin, 2007; Schroder et al., 2012; Yik et al., 2003). The CTDs of Fs(1)h and BRDT were also found to be capable of this interaction (Bisgrove et al., 2007).

BRDT is unique among the mammalian BET family proteins in that it is exclusively expressed in the testis (Jones, Numata, & Shimane, 1997); in mouse specifically in pachytene spermatocytes through to post-meiotic round spermatids (Shang, Nickerson, Wen, Wang, & Wolgemuth, 2007; Shang et al., 2004). BRDT is essential to support a normal gene expression program during meiosis and spermiogenesis. It was shown to be involved in the expression of over 3000 genes (Gaucher et al., 2012), two-thirds of which correspond to genes activated by BRDT and one-third of which are repressed by BRDT and correspond to genes mainly expressed in spermatogonia. In spermatogenic cells from 20 day-old testes, BRDT bound to acetylated transcription start sites (TSS) of genes that are highly expressed during meiosis and spermiogenesis, suggesting that BRDT has a direct role in modulating transcriptional activity in spermatogenic cells (Gaucher et al., 2012; Goudarzi et al., 2016). Moreover, in certain genes, BRDT mediates transcription by recruiting the P-TEFb complex to the TSS (Gaucher et al., 2012). Along with modulating transcription *per se*, BRDT also affects gene expression as part of splicing machinery and determining the length of the 3‟UTR of mRNA transcribed in round spermatids (Berkovits, Wang, Guarnieri, & Wolgemuth, 2012).

To understand the role of BRDT in regulating gene expression during spermatogenesis, we identified and characterized in detail the distribution and features of BRDT binding sites within gene units in the genome by performing genome-wide occupancy studies of BRDT complexes in enriched populations of pachytene spermatocytes and round spermatids. We found that BRDT preferentially binds to gene units of highly transcribed genes in spermatocytes and spermatids. Most of the BRDT binding sites co-localized with known histone modifications and histone variants that correlate with active transcription. Similarly, BRDT binding sites in promoters were enriched for the binding motifs of a several different transcription factors, including MYB, RFX, ETS and ELF1 in pachytene spermatocytes and JunD, c-Jun, CRE and RFX in round spermatids. This strongly suggests that BRDT has multiple binding partners, and that these different BRDT-containing complexes may have distinct functions that modulate the unique spermatogenic and spermiogenic transcriptional programs. Gene ontology analyses showed that the BRDT-bound genes corresponded to genes with essential roles in diverse cellular processes and in organelle functions that are essential for proper spermatogenesis and male fertility. Overall, these results define genomic features and signatures that characterize the BRDT binding sites in genes transcriptionally active in meiotic and post-meiotic cells, and strongly suggest transcriptional partners that may cooperate with BRDT to regulate transcriptional activity during spermatogenesis.

## Results and Discussion

### Overview of genome-wide analysis of BRDT binding in pachytene spermatocytes and round spermatids

As noted above, BRDT is expressed uniquely in the testis from early pachytene spermatocytes through round spermatids (Shang et al., 2004) and has been shown to be essential for spermatogenesis (Gaucher et al., 2012; Shang et al., 2007). In addition, previous ChIP-Seq analysis showed that while most of the BRDT binding sites are intergenic, a significant fraction of BRDT binds to acetylated TSS of genes activated by the end of meiosis (Gaucher et al., 2012). These observations lead us to explore in more detail the genomic binding sites of BRDT in enriched fractions of spermatocytes and spermatids, with a particular focus on gene units. We therefore used ChIP-Seq to map the genome-wide binding profiles of BRDT in formaldehyde-fixed chromatin isolated from samples enriched in pachytene spermatocytes and round spermatids using a C-terminal anti-BRDT antibody (Berkovits & Wolgemuth, 2011) and anti-IgG antibody as a control.

We identified 5452 BRDT-bound genomic loci in pachytene cell chromatin, of which 26% were located in promoters, 12% in exons, and 37% in introns, as designated in the annotated RefSeq genes (Figure 1A,B). The remaining 25% were in intergenic regions, defined as <u>not</u> in promoters, exons or introns. In round spermatids, we identified 5618 BRDT-bound genomic loci, of which 16% were located in promoters, 8% in exons, 43% in introns and 33% in intergenic regions (Figure 1A,B). These results differ somewhat from the BRDT distribution reported by Gaucher et al. (Gaucher et al., 2012), in that we found that 25% (spermatocytes) and 33% (round spermatids) of BRDT binding sites are intergenic, compared to their report of 66% and 60%, respectively. A possible explanation for the differences may be the criteria used to define and classify intergenic regions, which is key to avoid overlap between distinct regions, even more so when gene units are adjacent to one another. It should further be noted that different BRDT antibodies were used in each study. Studies on ChIP-Seq binding to intergenic regions of the other three BET proteins (BRD2, BRD3 and BRD4) in HEK293 cell lines were very similar to our findings (LeRoy et al., 2012). That is, ~65% of BRD-binding sites were located in genes (promoter or gene body) and ~35% of binding sites were intergenic. Although BRDT is the most divergent of the BET family proteins, and testicular cells are very different from HEK293 cells, the ratio of 2:1 gene binding to intergenic binding found by Leroy and colleagues more closely matches the ratios we observed for BRDT.

**Figure 1.**
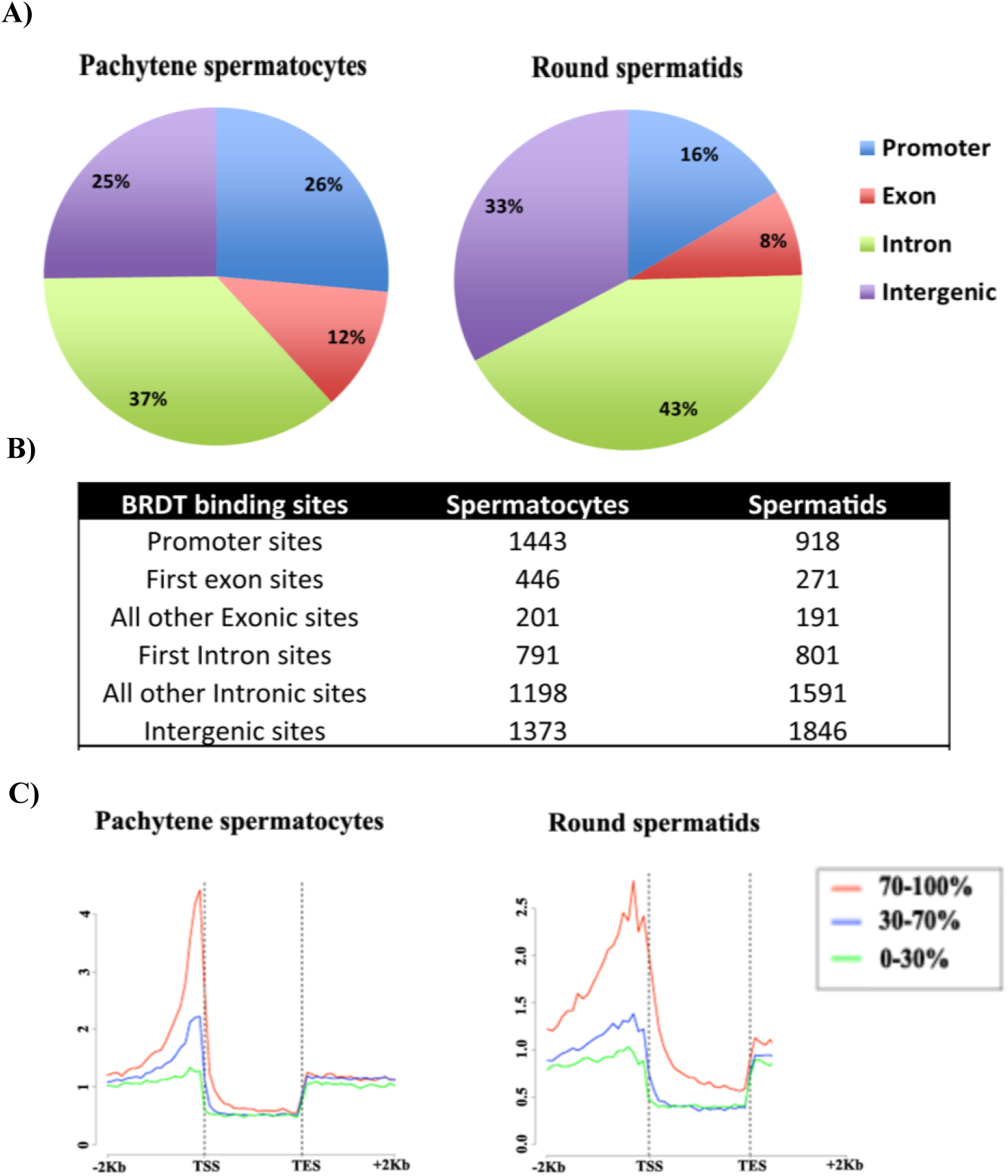
High–resolution genome-wide analysis of BRDT-containing complexes occupancy in pachytene spermatocytes and round spermatids of wild type mice (A-B) pie chart shows genomic annotation of location and number of BRDT binding sites across the genome in wild type pachytene spermatocytes and round spermatids. A promoter is defined as occupying 2 kb upstream of the transcription start site (TSS). (C) High-resolution analysis of average tag density across a gene body in wild type mouse testicular cells. In each plot, genes were classified into 3 groups based on expression levels in pachytene spermatocytes and round spermatids: highly expressed (the top 30% of genes indexed by expression levels), medium expression (the middle 40% of genes) and lowly expressed (the lowest 40% of genes). Average tag density across the gene unit was plotted for each group.

Considering that one of the most important functions of BRDT in spermatogenesis is to drive a developmentally regulated testis-specific gene expression program (Gaucher et al., 2012), we then focused our analysis on BRDT‟s role in transcription. We therefore associated BRDT binding sites with specific RefSeq genes, if the binding site was within 4 kb of the TSS. In spermatocytes, 1443 BRDT binding sites were located in the promoters of 1481 genes (Supplemental Table S1). This excess of genes above the number of binding sites is caused by the presence of a binding site between two genes that transcribe in opposite directions. That is, one binding site can be in the promoter of two different genes. In round spermatids, 918 BRDT binding sites were located in the promoters of 923 genes (Supplemental Table S2). Comparison of the two lists of putative BRDT target genes revealed that 225 genes bound BRDT in the promoter region in both cell types, which corresponded to 15% and 24% of the promoter-bound genes in spermatocytes and spermatids, respectively.

As three-quarters of the binding sites in spermatocytes and two-thirds of the sites in spermatids were within gene units (defined as the gene body plus promoter region), we analyzed the average BRDT tag density distribution across a gene. In both cell types, a major peak of BRDT occupancy was observed just 5‟ to the TSS (Figure 1C). Although a greater number of binding sites was present in the gene body as compared to the promoter, the promoter binding clustered one to two nucleosomes before the TSS, whereas the intronic and exonic binding sites were spread relatively evenly over the whole gene, albeit with a slight bias toward the 5‟ end of genes.

When all genes were indexed based on expression levels (as calculated from our earlier microarray expression analysis of pachytene spermatocytes and round spermatids) (Berkovits et al., 2012), the expression levels correlated positively with the degree of BRDT promoter occupancy. That is, the most highly expressed genes were most likely to have BRDT binding in the promoter (Figure 1C). In spermatocytes, 71% of the genes bound by BRDT in their promoters were highly expressed, 26% had medium expression levels and only 3% were expressed at low levels. In spermatids, the ratios were very similar, with 66% highly expressed, 30% with medium expression and 4% with low expression. These results are consistent with previously reported data showing the enrichment of BRDT binding sites at TSS +/− 1 kb and the direct correlation between genes binding BRDT at the TSS and their transcriptional activation in spermatocytes and spermatids (Gaucher et al., 2012).

### First intron and exon binding

We next investigated the distribution and characteristics of the BRDT binding sites in the gene bodies by first investigating BRDT‟s 5‟ intron and exon binding distribution. Introns in the 5′ proximal region of a gene („early‟ introns) have been shown to have important functional properties, often relating to the regulation of gene expression (Bradnam & Korf, 2008). The ability of an intron to act as an enhancer of gene expression has been termed intron-mediated enhancement (IME) (Mascarenhas, Mettler, Pierce, & Lowe, 1990). Further, the exons coding for 5‟ UTR can also be targets for transcriptional regulation (Al-Harthi & Roebuck, 1998; Turner et al., 2010). It is also possible that since the BRDT binding peaks were broad (usually over two nucleosomes), some might straddle the TSS. As we designated the midpoint of the peak as the BRDT binding site, some peaks might have been counted as being present in the first exon even though they partially overlap the promoter as well. Lastly, alternative promoter usage is prevalent in the testis (DeJong, 2006; Freiman, 2009; Liu, Zhang, Li, Yan, & Li, 2010) and some early intronic/exonic binding could in fact be binding of an alternative promoter. Interestingly, 791 of the 1989 (40%) intronic sites of BRDT binding in pachytene spermatocytes and 801 of the 2392 (33%) intronic sites of BRDT binding in round spermatids occur in the first intron (Figure 1B), the most common location for IME binding. Of the 647 exonic sites of BRDT binding in spermatocytes, 446 (69%) and 271 of the 462 (59%) exonic sites of BRDT binding in round spermatids occur in the first exon (Figure 1B).

### Enriched DNA binding sites in BRDT-bound regions suggest possible interaction with transcription factors

We next asked whether the promoter regions bound by BRDT would be enriched with distinct transcription factors (TFs) binding motifs, which would suggest a possible interaction between BRDT and specific TFs. Consensus and optimal *in vitro* DNA binding motifs in BRDT-bound regions were therefore ascertained from JASPAR, UniPROBE and Transfac. MEME was used to find motifs enriched in the 500 BRDT promoter-binding peaks with the highest number of reads in each cell type. Each motif was scored for significantly increased binding frequency at these peaks as compared to background. In pachytene spermatocytes, the four most highly enriched motifs were for binding of the TFs MYB, RFX, ETS and ELF1 (Figure 2A) and in round spermatids, the four highest peaks were for JunD, c-Jun, CRE and RFX (Figure 2A). This is of particular interest as it suggests a dynamic interaction between TFs and epigenetic readers such as BRDT, which could be recruited to promoters to co-regulate a subset of genes during spermatogenesis.

**Figure 2.**
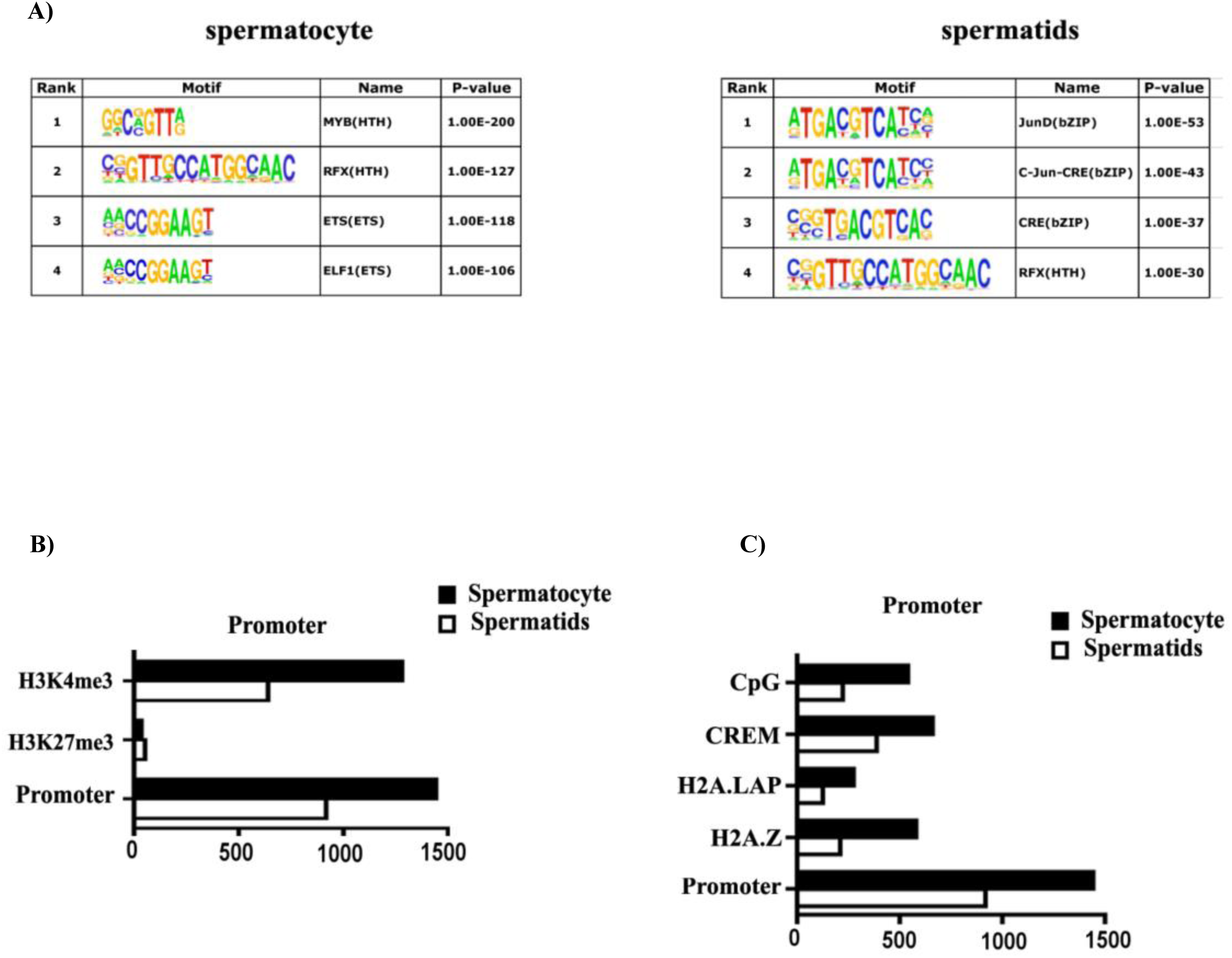
Characterization of BRDT binding site occupancy. (A) BRDT in vivo consensus binding site predicated by the MEME and presented as a sequence LOGO. A total 500 peak regions were analyzed for overrepresented motif. Enriched transcription factor motifs lying within BRDT binding sites in pachytene spermatocyte and round spermatids of wild type mice. (B) Correlative analysis of H3K4me3, H3K27me3 with BRDT binding site occupancy. (C) Correlative analysis of CpG, CREM, H2A.LAP, H2AZ with BRDT binding site occupancy. The x-axis in (B) and (C) refers to the number of associations with BRDT binding sites.

Our MEME analysis revealed that MYB motifs were the most prevalent in spermatocytes (p-value 1.0 E-200). MYB proteins are known transcriptional regulators in spermatocytes, particularly A-MYB, which is a master regulator of meiotic genes involved in multiple processes in spermatocytes (Bolcun-Filas et al., 2011). A-MYB has an expression pattern in spermatocytes similar to that of BRDT and depletion of either protein results in arrest at the pachytene stage (Gaucher et al., 2012; Latham et al., 1996; Shang et al., 2007; Toscani et al., 1997). Additionally, A-MYB has been shown to bind and up-regulate the *Rfx2* promoter and conversely RFX2 has been suggested to be a candidate downstream amplifier of A-MYB regulation (Horvath, Kistler, & Kistler, 2009; Toscani et al., 1997). Interestingly, RFX motifs were also enriched at BRDT binding sites in promoters in both cell types examined. While several RFX family TFs are up-regulated in post-meiotic cells, RFX2 alone is expressed at high levels in pachytene spermatocytes (Horvath et al., 2009). Of note, the promoter of the testis-specific histone 1 variant *H1t*, which we had previously shown to be a direct target of BRDT-containing complexes (Shang et al., 2007), contains two imperfect inverted repeats which together comprise the X-box motif of RFX binding (VanWert, Wolfe, & Grimes, 2008). Of the 225 genes that were promoter-bound by BRDT in both spermatocytes and spermatids, 75% of them contained the RFX motif in their promoters.

Our results further show that the TFs that putatively associate with BRDT are specific to the development stage of the germ cells. For example, in round spermatids, the highest scores were observed for JunD, C-Jun, and CRE (Figure 2A), consistent with the higher level of expression of *junD* mRNA in post-meiotic round spermatids than in meiotic pachytene spermatocytes (Alcivar, Hake, Kwon, & Hecht, 1991) and suggesting this could influence its potential specific interaction with BRDT in these cells. Also, JunD-deficient males exhibited multiple age-dependent defects in reproduction and impaired spermatogenesis with abnormalities in sperm head and flagellum structures (Thepot et al., 2000) that are similar to *BRDT*^Δ*BD1*/Δ*BD1*^ mutant mice (Shang et al., 2007).

Overall, these observations suggest that BRDT may bind to specific TFs in distinct cellular contexts, and diverse BRDT containing-complexes may be recruited to different promoters to regulate specific genes throughout spermatogenesis.

### The majority of BRDT binding sites in gene units co-localize with active chromatin histone marks

To further assess the association of BRDT promoter binding with active transcription and the epigenetic features of the BRDT binding sites at promoter regions, we next compared our BRDT ChIP-Seq data with published ChIP-seq data sets for features and markers of active transcription.

The active chromatin histone mark H3K4me3 and the repressive histone mark H3K27me3 have been examined in extracts obtained from total testicular cells (Cui et al., 2012). Comparison of these data with our results for BRDT binding revealed that ~89% (1279) and ~70% (639) of BRDT promoter binding sites are associated with H3K4me3 in pachytene spermatocytes and round spermatids respectively, and only 2.4% (52) and 5.7% (34) of BRDT promoter binding sites associated with H3K27me3 in the two populations (Figure 2B).

We further extended this analysis of BRDT binding and marks of active transcription by comparing our data with published ChIP-Seq data sets for CpG islands and the histone H2A variants H2A.Z and H2A.LAP1, which had been examined in extracts obtained from total testicular cells (Cui et al., 2012; Martianov et al., 2010; Soboleva et al., 2011). The location of CpG islands and GC-enriched regions to transcriptionally permissive chromatin is well-documented (R. Illingworth et al., 2008; R. S. Illingworth & Bird, 2009), although the role of the CpG island as a sequence motif in chromatin organization and, consequently, in gene regulation, is not yet clearly understood. During spermiogenesis it is known that transition protein 2 (TP2) can use CpG island sequences to reorganize spermatid chromatin (Kundu & Rao, 1996). In our study, we found that 37.4% (540) and 24.3% (223) of BRDT promoter binding sites in pachytene spermatocytes and round spermatids, respectively, overlapped CpG islands (Figure 2C). Both of the histone H2A variants H2A.Z and H2A.LAP1 have been reported to be components of the chromatin of active promoters (Soboleva, Nekrasov, Ryan, & Tremethick, 2014). Interestingly, it has been reported that BRD2 is recruited to androgen receptor (AR)–regulated genes in an H2A.Z-dependent manner and that chemical inhibition of BRD2 recruitment greatly inhibits AR– regulated gene expression (Draker et al., 2012). We found that 40% (580) and 23% (212) of BRDT promoter binding sites associated with H2A.Z in pachytene spermatocytes and round spermatids, respectively (Figure 2C). The association of BRDT promoter binding sites with H2A.LAP1, however, revealed overlap for only 19.2% (276) and 13.6% (125) of binding sites in pachytene spermatocytes and round spermatids, respectively (Figure 2C).

Our MEME analysis mentioned above identified CRE motif binding sites in spermatids (Figure 2A) and it is known that the CREMτ isoform of the TF CREM is strongly and selectively expressed in round spermatids (Blendy, Kaestner, Weinbauer, Nieschlag, & Schutz, 1996; Foulkes, Mellstrom, Benusiglio, & Sassone-Corsi, 1992), suggesting that BRDT is associated with CREMτ to potentially regulate gene expression in these cells. To this end, we next compared our BRDT ChIP-Seq data with CREMτ ChIP-Seq data (Martianov et al., 2010). Our analysis revealed a notable overlap of BRDT and CREMτ binding in promoters, with 42% (389) of all BRDT binding sites shared with reported CREMτ binding sites (Figure 2C). This overlap indicates that a possible collaboration between these two factors during spermiogenesis. For example, CREB binding protein (CBP), which binds to CRE elements, can catalyze lysine butyrylation in histones (Chen et al., 2007), and butyrylated H4K5 is sufficient to abolish the interaction between BRDT and histone H4 (Goudarzi et al., 2016). Thus, the presence of BRDT and other histone modifiers in the same genomic regions could allow an interplay that produces a negative feedback regulatory loop, releasing BRDT from promoter complexes.

### GO enrichment analysis of BRDT-bound at the promoters of genes in spermatocytes and spermatids

As BRDT is a known transcriptional modulator and our results suggest that BRDT could be interacting with TFs in a developmental stage-specific epigenetic pattern, we next explored the possible physiological relevance in spermatogenesis for those genes that had BRDT binding in their promoters (Supplemental Table S1 and S2). To obtain a global overview of the biological functions of proteins encoded by BRDT-bound genes at the promoters, we performed a functional annotation using DAVID (Database for Annotation, Visualization, and Integrated Discovery) analysis for biological processes Gene Ontology (GO) terms (Huang da, Sherman, & Lempicki, 2009a, 2009b) (Supplemental Table S3 and S4). As we found that BRDT promoter-binding was robust at genes that are highly transcribed in germ cells, we predicted that BRDT would be bound to promoters of genes with important functions during spermatogenesis. GO enrichment analysis revealed that in spermatocytes, the BRDT-bound gene set was most significantly enriched for RNA processing (GO:0006396; *P*-value = 1.79E-15) which includes mRNA metabolic process (GO:0016071), followed by chromosome organization (GO:0051276; *P*-value = 2.74E-15), and cell cycle process (GO:0022402; *P*-value = 1.39E-14) (Table 1A; Supplemental Table S5). The GO enrichment analysis suggests that beyond the biological processes previously associated with BRDT, such as RNA processing, chromosome organization, and cell cycle process, we observed enrichment for genes involved in biological processes not clearly yet associated to BRDT function, including protein ubiquitination, ncRNA metabolic process, and DNA-templated transcription, initiation (Table 1A; Supplemental Table S5).

**Table 1.** GO-terms for enriched biological processes in our BRDT-bound genes at the promoters in (A) spermatocytes and (B) spermatids.

| **A) GO-term biological process (Spermatocytes)** | **pval** |
| --- | --- |
| GO:0006396~RNA processing | 1.79E-15 |
| GO:0051276~chromosome organization | 2.74E-15 |
| GO:0022402~cell cycle process | 1.39E-14 |
| GO:0044085~cellular component biogenesis | 1.75E-14 |
| GO:0016567~protein ubiquitination | 8.50E-14 |
| GO:0006259~DNA metabolic process | 2.22E-13 |
| GO:0044257~cellular protein catabolic process | 2.24E-12 |
| GO:0034660~ncRNA metabolic process | 2.90E-10 |
| GO:0033365~protein localization to organelle | 2.58E-09 |
| GO:0006352~DNA-templated transcription, initiation | 1.86E-08 |
| **B) GO-term biological process (Spermatids)** |  |
| GO:0007283~spermatogenesis | 7.36E-15 |
| GO:0070925~organelle assembly | 3.24E-08 |
| GO:0016071~mRNA metabolic process | 3.13E-07 |
| GO:0016567~protein ubiquitination | 3.57E-07 |
| GO:0044257~cellular protein catabolic process | 9.73E-07 |
| GO:0033365~protein localization to organelle | 2.04E-06 |
| GO:0044085~cellular component biogenesis | 1.11E-05 |
| GO:0051168~nuclear export | 1.56E-05 |
| GO:0007017~microtubule-based process | 1.89E-05 |
| GO:0006470~protein dephosphorylation | 2.41E-05 |

In contrast, in spermatids, spermatogenesis (GO:0007283; *P*-value = 7.36E-15) was the most significantly enriched for biological process, followed by organelle assembly (GO:0070925; *P*-value = 3.24E-08), and mRNA metabolic process (GO:0016071; *P*-value = 3.13E-07) (Table 1B; Supplemental Table S5). Beyond the biological processes previously associated with BRDT, such as spermatogenesis, mRNA metabolic process and protein localization to organelle, we observed enrichment for genes involved in biological processes not yet clearly related to BRDT function, including protein ubiquitination, microtubule-based process and protein dephosphorylation (Table 1B; Supplemental Table S5).

Taking into consideration the overall content of the two cell types, the overlap of 5 biological processes in both spermatocytes and spermatids is highly statistically significant; RNA processing/mRNA metabolic process, cellular component biogenesis, protein ubiquitination, cellular protein catabolic process and protein localization to organelle (Table 1). We note that mRNA metabolic process (GO:0016071) which is significantly enriched GO-term in spermatids is a derivative function of RNA processing (GO:0006396) which is significantly enriched GO-term in spermatocytes (according to the GO annotation). Our lab has previously investigated the function of BRDT in mRNA processing which includes 3′-UTR processing and alternative splicing, thereby affecting gene and protein expression (Berkovits et al., 2012). Based on our findings, BRDT was suggested to function in modulating gene expression as part of the splicing machinery and may partially be explained by occupancy of BRDT at promoters of RNA processing/mRNA metabolic process genes. These genes included *Hnrnph1, SNRPD3, RBM7, CELF6, RBM39*, among others (Supplemental Table S5).

Protein ubiquitination was also significantly enriched for biological process in both two cell types. Although BRDT has not been yet to related to protein ubiquitination pathway, this is of interest in light of the previously studied functions of ubiquitination-related genes in spermatogenesis (Bose, Manku, Culty, & Wing, 2014). Our BRDT ChIP-seq analysis identified 93 and 50 genes in spermatocytes and spermatids, respectively, that are related to the ubiquitin pathways (Table 1;Supplemental Table S5). Genes bound by BRDT encode E2 ligases, HECT type E3 ligases, U-box type E3 ligases, single-Ring finger E3 ligases, components of a multi-subunit Cullin-Rbx type E3 ligase complexes, APC/C E3 ligases and *ubiquitin B* (*Ubb*) itself (Supplemental Table S5). It has been shown that the mouse *Ubb* is essential for meiotic progression: *Ubb-/-* mice of both genders are infertile due to the cell death during pachytene, and males exhibit complete testicular degeneration by 2 years of age (Ryu et al., 2008). Another example is *ubiquitin-conjugating enzyme E2B* (*Ube2b*), a histone E2 ligase that is required for fertility and effects proper gene expression and chromatin condensation in spermatids (Mulugeta Achame et al., 2010; Roest et al., 1996), phenotypes that are similar to those seen in *BRDT*^Δ*BD1*/Δ*BD1*^ mutant mice (Gaucher et al., 2012; Shang et al., 2007).

### Visualization of the localization of BRDT-binding sites at promoters

Based on the enriched biological processes identified in our GO analysis and potential relevance to BRDT‟s functions as shown in previous studies from our laboratory (Berkovits et al., 2012; Berkovits & Wolgemuth, 2011; Manterola et al., 2018; Wang & Wolgemuth, 2016), we selected three critical processes and specific examples with which to localize the sites of BRDT binding, namely RNA processing/mRNA metabolic process, chromosome organization and spermatogenesis (Figure 3). In spermatocytes and spermatids, BRDT bound at the promoter of 106 and 43 genes, respectively, that are related to RNA processing and mRNA metabolic process, respectively (Supplemental Table S5). Genome browser visualization of the BRDT-bound at the promoter of gene loci in both cell types showed a distinct patterns of BRDT binding to promoters between the cell types: both spermatocytes and spermatids, only spermatocytes and only spermatids. Genes that are major components of the spliceosome and directly affect alternative splicing, for example *Hnrnph1* (Masuda et al., 2008), are bound by BRDT in both cell types (Figure 3A, left). BRDT binding at this class of genes is not surprising as we have shown that partial loss of BRDT function has an effect on mRNA splicing in both spermatocytes and round spermatids (Berkovits et al., 2012). *Ppp4r2*, protein phosphatase 4 regulatory subunit 2 which plays a role in processing of spliceosomal snRNPs (Carnegie et al., 2003), is bound by BRDT only in spermatids (Figure 3A, middle). Interestingly, we have shown that *Ppp1r8*, protein phosphatase 1 regulatory subunit 8, which is involved in pre-mRNA splicing, has a loss of truncated 3′-UTR in *Brdt* mutant round spermatids (Berkovits et al., 2012). Cleavage stimulation factor, 3’ pre-RNA subunit 2, *Cstf2t*, which plays a significant role in mRNA polyadenylation and splicing of *Cyclic AMP-Responsive Element Modulator (CREM)* in mouse testis (Grozdanov, Amatullah, Graber, & MacDonald, 2016) is bound by BRDT only in spermatocytes (Figure 3A, right). Male mice that lack *Cstf2t* (*Cstf2t-/-* mice) experience disruption of spermatogenesis and are infertile (Li et al., 2012).

**Figure 3.**
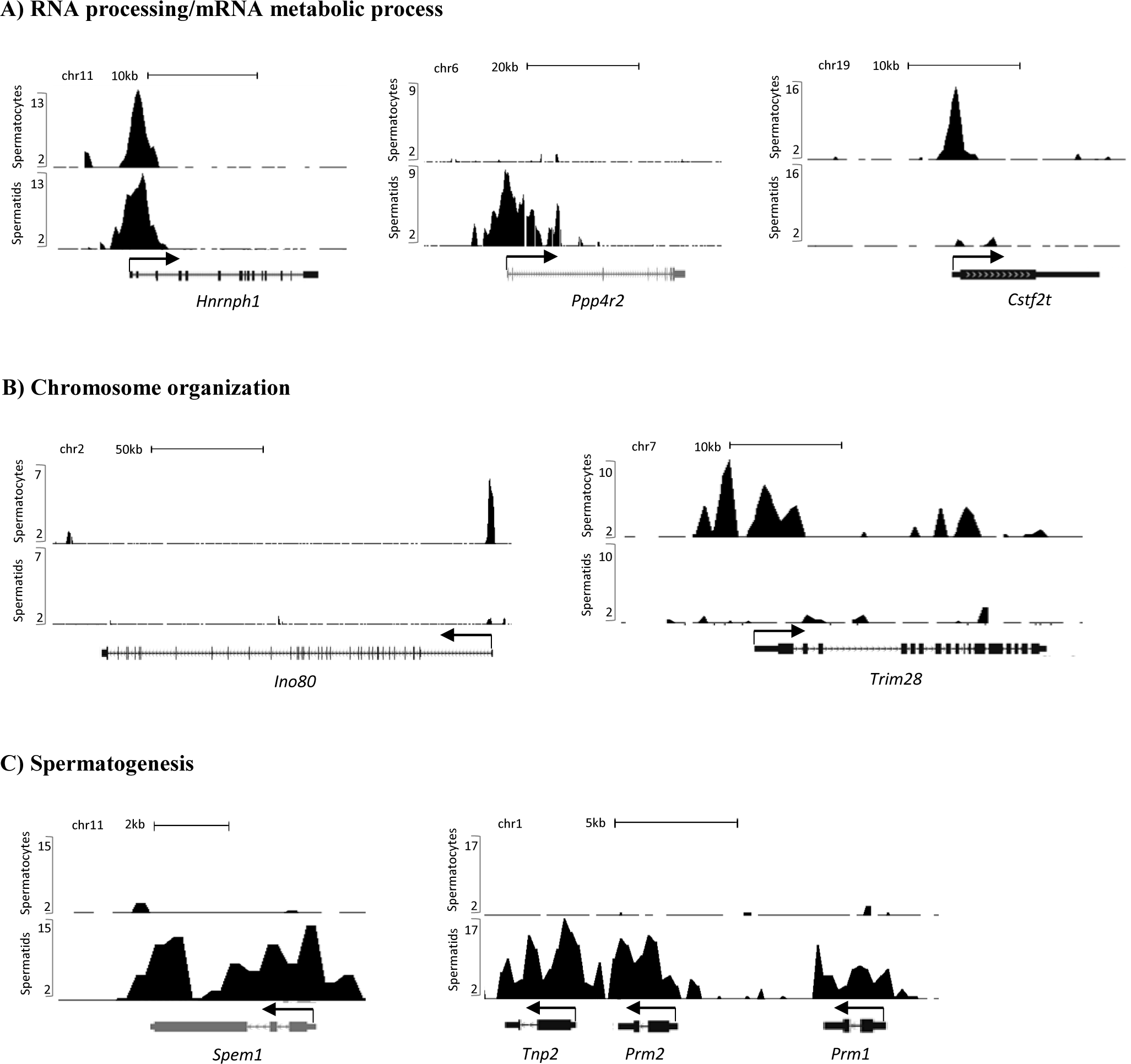
UCSC Genome Browser screenshot of BRDT ChIP-seq signals at the gene promoters implicated in (A) RNA processing/mRNA metabolic process, (B) chromosome organization and (C) spermatogenesis. Graphic representation of ChIP-Seq reads derived from pachytene spermatocytes and round spermatids on all panels. ChIP-seq signals are displayed as normalized RPM (reads per million reads) values.

In terms of the chromosome organization biological process, which is most secondly enriched for biological process in spermatocytes, *Ino80* and *Trim28* both showed a strong signal peak of BRDT. *Ino80* is a catalytic ATPase subunit of the INO80 chromatin remodeling complex which is involved in transcriptional regulation, DNA replication and DNA repair (Hur et al., 2010; Jiang et al., 2010; Jin et al., 2005). During spermatogenesis, INO80 is expressed in developing spermatocytes during the early stages of meiotic prophase I, and loss of INO80 (*Ino80*^*cko*^) results in an arrest during meiosis associated with a failure to repair DNA damage during meiotic recombination (Serber, Runge, Menon, & Magnuson, 2016). Our ChIP-seq analysis showed that BRDT bound its promoter in spermatocytes (Figure 3B, left). As another example of genes involved in chromosome organization, TRIM28 mediates gene silencing by recruiting various heterochromatin-inducing factors, such as heterochromatin binding protein (HP1 α, β, γ), the histone methyltransferase SETB1 and the nucleosome remodeling and histone deacetylation (NuRD) complex (Schultz, Ayyanathan, Negorev, Maul, & Rauscher, 2002; Schultz, Friedman, & Rauscher, 2001; Sripathy, Stevens, & Schultz, 2006). During spermatogenesis, TRIM28 is expressed in pachytene and diplotene spermatocytes and round spermatids and its ablation produces testicular degeneration due to shedding of these cells (Weber et al., 2002). We have shown that TRIM28 interacts with BRDT and that this complex binds to the *H1t* promoter and modulates its transcription (Wang & Wolgemuth, 2016). The binding of TRIM28-BRDT to *H1t* was lost in *BRDT*^Δ*BD1*/Δ*BD1*^ mutant mice and concomitantly, there were elevated levels of *H1t* expression and reduced levels of the repressive histone mark H4R3me2 in spermatids (Shang et al., 2007; Wang & Wolgemuth, 2016). Interestingly, our ChIP-seq analysis revealed that the *Trim28* promoter is bound by BRDT exclusively in spermatocytes (Figure 3B, right), suggesting that BRDT may be involved in TRIM28-mediated chromosome organization and transcription repression in spermatids by modulating *Trim28*‟s expression in pachytene spermatocytes.

In spermatids, spermatogenesis was the most significantly enriched GO term for biological process, and included several genes known to be essential for male fertility. *SPEM1*, for example, is required for proper removal of the cytoplasm from the spermatid head, and also has a role in nucleocytoplasmic transport and intra-manchette transport (Bao et al., 2011; Bao, Zhang, Zheng, Xu, & Yan, 2010; Zheng et al., 2007). *Spem1* is bound by BRDT extensively in spermatids throughout its genomic region (Figure 3C, left). Strong BRDT binding was also observed in spermatids not only at the promoter but also throughout the genomic region. Also in the spermatogenesis GO biological process are the sperm-specific nuclear proteins *protamine 1* and *2* (*Prm1*, *Prm2*) and *transition protein 2* (*Tnp2*), which are essential in generating elongating spermatids and mature spermatozoa (Brewer, Corzett, & Balhorn, 1999), and are involved in intermediate stages in the replacement of histones by protamines (Meistrich, Mohapatra, Shirley, & Zhao, 2003; O’Carroll et al., 2000; Yu et al., 2000) (Figure 3C, right).

In summary, to further elucidate the BRDT function in spermatogenesis, we performed ChIP-Seq to examine in detail the genomic features of BRDT bound chromatin in spermatocytes and spermatids. The results presented here show that the majority of BRDT binding sites were located in genes (the promoter or gene body), and that the binding sites co-localize with active chromatin marks. Further, we identified enrichment for specific TF motifs, such as MYB, RFX, ETS and ELF1 motifs in pachytene spermatocytes and JunD, c-Jun, CRE and RFX motifs in round spermatids. We also found that BRDT binds to the gene body of highly transcribed genes in spermatocytes and spermatids, which by gene ontology analysis, correspond to genes with essential roles in diverse biological processes that are essential for proper spermatogenesis and male fertility. The GO analysis also revealed that there are unique sets of genes enriched in biological processes between the two cell types, suggesting distinct developmentally stage-specific functions for BRDT. Our ChIP-Seq analysis of the BRDT–bound genomic loci in pachytene spermatocytes and round spermatids thus provides a valuable resource for understanding the function of BRDT in male germ cell differentiation.

## Materials and Methods

### Mice

The mice used in the present analysis were maintained in a National Institute of Health Research Animal Accredited Facility in accordance with the specifications of the Association for Assessment and Accreditation of Laboratory Animal Care. Mouse protocols were approved by the Institutional Animal Care and Use Committee of Columbia University.

### Generation of populations of pachytene spermatocytes and round spermatids

The pachytene spermatocyte sample consisted of cellular suspensions obtained from testicular tubules isolated from eighteen day old male 129 Sv/Ev mice following our laboratory’s standard protocols as previously published (Wolgemuth, Gizang-Ginsberg, Engelmyer, Gavin, & Ponzetto, 1985). Briefly, testes were decapsulated and transferred to cold PBS, the seminiferous tubules were manually sheared with scissors and by pipetting, and then passed through a 40 μm filter. Although this population also includes spermatogonia, leptotene/zygotene spermatocytes, and some Sertoli cells, these cells do not express BRDT. Round spermatids were purified as previously described (Wolgemuth et al., 1985). Briefly, testes were dissected from seven adult mice and decapsulated. The cellular suspension was obtained from the tubular mass by digestion with 1.0 mg/ml collagenase (Sigma Cat# C0130) in RPMI buffer followed by digestion with 0.25 mg/ml trypsin (Sigma cat#T8003) in RPMI (Wolgemuth et al., 1985). Cell aggregates and connective tissue were removed by filtration through 74-pm nylon mesh (BD Falcon). The single cell suspension of germ cells was separated using gravity sedimentation on a 2-4% BSA in DPBS gradient. Fractions containing round spermatids were pooled to yield samples containing ≥90% round spermatids, as assessed by flow cytometric analysis (Golan et al., 2003) on a BD FACSCalibur Cell Analyzer. All steps of cell separation, fractionation and pooling were performed at 4°C in order to maintain cell viability and maximize nucleic acid integrity.

### Chromatin immunoprecipitation coupled to massively parallel sequencing (ChIP-Seq)

To identify BRDT-containing complex binding sites, we prepared cross-linked chromatin samples for ChIP-Seq analysis. Approximately 10^7^ cells from the two cellular preparations were fixed with 1% formaldehyde for 10 min at room temperature. The chromatin template was then fragmented to a size of 150 to 200 bp by sonication (Cuddapah et al., 2009). Aliquots of the chromatin samples were subjected to chromatin immunoprecipitation using a C-terminal anti-BRDT antibody generated by our laboratory (Shang et al., 2007) or anti-IgG as a control. This was followed by performing end repair and addition of “A” bases to the 3‟ end of the DNA fragments, and ligation of adapters to the DNA fragments (Cuddapah et al., 2009). The resulting fragments were then size-selected for 150-350bp regions before generating the library, which was enriched for the adapter-modified DNA fragments by PCR. The purified DNA samples were subjected to analysis on the Ilumina Cluster Station and Genome Analyzer. In samples using the BRDT antibody, 17.5 and 18 million uniquely aligned short sequence reads in pachytene spermatocytes and round spermatids, respectively, were obtained. Using the IgG antibody, 13 and 8.3 million reads in the two populations were obtained.

### Data preprocessing

Sequenced tags were mapped to the mouse genome (mm8) using Bowtie algorithm (Langmead, Trapnell, Pop, & Salzberg, 2009) with the option „-m 1‟. In order to reduce PCR amplification bias, at most one read was kept per genomic location. The resulting lists of uniquely mapped non-redundant reads were used for all downstream analyses. UCSC genome browser displayable BedGraph files for ChIP-Seq data were generated by counting tags in 200-bp windows across the genome and normalizing window tag counts by 10 million total sequenced reads to make different libraries directly comparable.

### Genome-wide distribution of binding sites

BRDT binding sites from the ChIP-Seq data were identified using SICER software (Zang et al., 2009) using IgG data as control and a False Discovery Rate (FDR) threshold of 10%. The midpoint of each identified BRDT binding region („island‟) is regarded as a BRDT binding site. Gene and exon location information was obtained from the Ensembl database (www.ensembl.org) using the BioMart tool. Based on this information, promoter, exonic, intronic, and intergenic regions were defined as follows. A promoter was defined as being 2 kb upstream of the TSS. To avoid double counting at the promoter overlaps, coordinates of all promoters were merged to define a non-redundant set of promoter regions. Similarly, other classes of regions (exons and introns) were merged to avoid double counting. Some BRDT binding sites are located in multiple classes of region, e.g. in the exon of one transcript isoform and in the intron of another transcript isoform of the same gene. In order to deal with such ambiguities in the assignment of BRDT binding sites to different classes, we used the following prioritization: promoter took precedence over an exon, followed by exon > intron > intergenic. Thus, for example, if a BRDT binding site is located in both an exon and an intron, then it is assigned as being located in the exon. The number of BRDT binding sites in the resulting non-redundant and non-overlapping classes of regions were counted and used to create pie charts to illustrate the distribution of binding sites.

### Gene body assay

The sequence read density (per 100-bp window and 30 million non-redundant uniquely mapped reads) was determined along the gene-body and in the upstream/downstream 2 kb regions in the sample and control IgG libraries. The signal level in each window was computed as the difference between the read densities in the sample and control libraries, with the negative values set to zero. Genes were ranked based on their expression levels and stratified into three expression groups: the „high‟ group (the top 30% genes in the ranked list), the „silent‟ group (the bottom 30% genes), and the „medium‟ group (the remaining genes). The mean signal levels over all genes in each group were determined and displayed as gene-body plots.

### Motif analysis

MEME (Multiple Expectation maximization for Motif Elicitation) (Bailey et al., 2009) with default parameters was used to identify statistically overrepresented motifs within the inferred binding sites. In order to reduce noise for the motif identification, we used only DNA sequences at the top 500 BRDT binding sites in terms of ChIP-Seq tag density.

### Comparison of our ChIP-seq data with the published ChIP-seq data on CREM, and H2A-variant, H3K4me3 and H3K27me3

As defined above, BRDT binding sites are midpoints of BRDT binding regions identified by the SICER algorithm. To assess the overlap of BRDT binding with regions previously reported to be bound by selected nuclear proteins, including CREM (Martianov et al., 2010), the H2A variants H2A.Lap1 and H2AZ (Nekrasov et al., 2012), and H3K4me3 and H3K27me3 (Cui et al., 2012), the chromosomal coordinate of the published binding regions of these proteins was designated as (x1, x2). A BRDT binding site x is assumed to overlap this region if x1 ≤ x ≤ x2.

### Functional enrichment analysis

DAVID Bioinformatics Resources 6.8 was used to analyze the gene lists derived from ChIP-seq data to find enriched biological functions (Huang da et al., 2009a, 2009b). We uploaded the significant BRDT-bound at the promoters of genes to investigate the enriched potential functions. *P*-value < 0.1 and false discovery rate (FDR) < 0.05 were used as the cut-off criteria for the analysis.

## Availability of data and materials

The GEO accession number for the raw and processed ChIP-Seq data described in this article is GSE98489.

## Acknowledgments

This work was supported in part by a grant from the NIH, GM081767 (D.J.W), by the Division of Intramural Research, NHLBI, NIH (K.Z.), and an “Estadías Cortas de Investigación Internacionales”-Internationalization Grant UCH-1566 (M.M).

